# Harnessing subtractive genomics for drug target identification in *Streptococcus agalactiae* serotype v (atcc baa-611 / 2603 v/r) strain: An *in-silico* approach

**DOI:** 10.1101/2025.02.04.636418

**Authors:** Ashiqur Rahman Khan Chowdhury, Farjana Yasmin Tithi, Nusrat Zahan Bhuiyan, Afsana Ferdousi Ishita, Md Mahmodul Hasan Sohel

## Abstract

Developing a therapeutic target for bacterial disease is challenging. *In silico* subtractive genomics methodology offer a promising alternative to traditional drug discovery methods. *Streptococcus agalactiae* infections depend on two crucial criteria: drug-resistance and the existence of virulence factors. It is essential to underline that *S. agalactiae* strains have emerged to be resistant to several drugs. Hence, there is a need for research on novel drugs and techniques that are potent, economical, productive, and dependable to combat *S. agalactiae* infections. In this study advanced computational techniques were exploited to examine potential druggable targets exclusive to this pathogen. Our study uncovered 200 non-homologous proteins in *S. agalactiae* serotype V (Strain ATCC BAA-611 / 2603 V/R) and identified 68 essential proteins indispensable for the bacterium’s survival. Therefore, these 68 proteins are potential targets for drug development. Subcellular localization analysis unveiled that the pathogen’s cytoplasmic membrane contained essential proteins among these vital non-homologous proteins. On the other hand, based on virulent protein predictions, six proteins were seen to be virulent. Among these, we prioritized two proteins (Sensor protein LytS and Galactosyl transferase CpsE which are exclusively found in *S. agalactiae*) as potential druggable targets and selected them for further structural investigation. The proteins chosen could serve as a foundation for the identification of a promising therapeutic compound that has the potential to neutralize these enzymatic proteins, thereby contributing to the reduction of risks linked to the drug-resistant *S. agalactiae*.

## Introduction

*S. agalactiae*, is a gram-positive, β-haemolytic coccus, commonly known as a group B streptococcus or GBS. This encapsulated facultative anaerobe is catalase-negative and features a group B antigen in its thick peptidoglycan cell wall, with a tendency to form chains [1,2]. It is commonly found as a normal part of the body’s flora in the urogenital and lower gastrointestinal tracts, and may also be present in the oropharynx [3–7]. Based on the composition of the capsular polysaccharide, a key virulence factor, this bacterium is classified into 10 serotypes (Ia, Ib, II-IX) [1,8–10].

In 1935, Rebecca Lancefield was the first to describe the colonization of the vagina by *S. agalactiae* and the first description of the bacterium as a human pathogen and its role in causing invasive illness was documented in 1964 [11,12]. It is mostly responsible for multiple infections including sepsis, meningitis, soft-tissue infection, urinary tract infection, pneumonia, and premature delivery [2,13–15]. Newborns of 24-48 hours old are more susceptible (approximately 33.3%) to this pathogen compared to those of older than 48 hours (only 8%) [16]. Among adults, this bacterium has the potential to induce peripartum chorioamnionitis and bacteremia in new mothers (occurring in 0.03% of cases) as well as endocarditis and osteomyelitis in other individuals [2,17]. Individuals who are immunocompromised, elderly, or have conditions such as diabetes mellitus, alcoholism, cancer, cirrhosis, a history of stroke, or HIV/AIDS are at increased risk of Group B streptococcal infection [2,17,18]. The pathogen continues to be a primary cause of newborn sepsis, resulting in around 90,000 fatalities in early infancy and at least 57,000 stillbirths worldwide [19]. This bacterium is becoming increasingly resistant to traditional drugs, therefore, to effectively prevent and treat infections, it is vital to find out novel drug targets for *S. agalactiae* serotype V.

The exceptional advancements in computational biotechnology and bioinformatics have significantly impacted drug design, leading to reduced expense and duration for traditional laboratory trials by facilitating the discovery of therapeutic candidates, structure-based drug development, and the identification of host-specific targets through genomic data analysis. A key focus has been on the subtractive genomics technique, which aims to identify proteins with therapeutic potential exclusive to pathogenic genomes while excluding similar host proteins. This methodology has successfully identified novel species-specific therapeutic targets across various pathogenic strains including [20–39].

In this study, we aim to develop a potential therapeutic target to combat *S. agalactiae* serotype V through the application of sophisticated computational biotechnology and bioinformatics approaches, addressing the pressing global issue of drug resistance. Thus, in order to analyze the proteome of *S. agalactiae* serotype V in its entirety, the subtractive genomics technique was applied. Essential proteins for pathogen’s survival were prioritized using computational tools, followed by the removal of host homologous proteins to minimize therapeutic interference. The remaining pathogenic proteins were examined for subcellular localization and virulent capabilities, leading to the recognition of cytoplasmic membrane proteins and their virulence potential. Two promising proteins, Sensor protein LytS and Galactosyl transferase CpsE, were identified as potential therapeutic targets. These enzymatic proteins underwent further protein network analysis and structural investigation, highlighting that they could potentially have the promise to serve as prospective selections for the development of vaccines or drugs that specifically target *S. agalactiae* serotype V.

## Materials and Methods

The comprehensive protocol for discovering proteins that are essential and unique to *S. agalactiae* serotype V (Strain ATCC BAA-611/2603V/R) for potential drug target identification is outlined in **Fig 1**.

**Fig 1.**
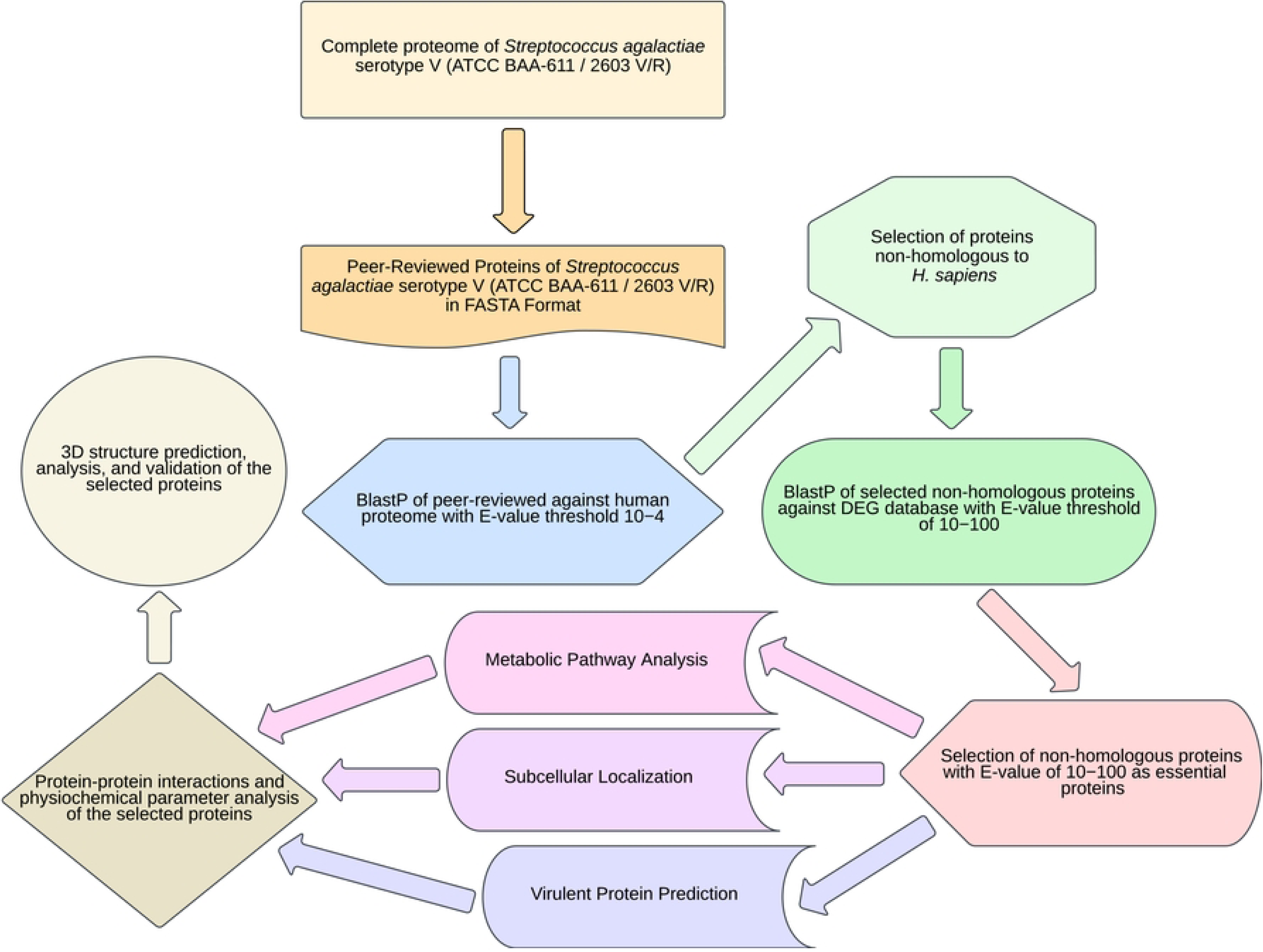
Flowchart for identifying potential drug targets in *S. agalactiae* serotype V. This shows the step-by-step process for identifying drug targets through the subtractive genomics approach.

### Protein Sequence Retrieval

The whole proteome of *S. agalactiae* serotype V (Strain ATCC BAA-611/2603V/R) was retrieved from UniProt database [40–42] along with its peer-reviewed proteins in FASTA format.

### Finding Non-homologous Proteins

To distinguish proteins that are non-homologous to the host, a BLASTp search of the NCBI database (with e-value threshold of 10^−4^) was conducted against *Homo sapiens* on the selected peer-reviewed proteins [43]. The non-homologous protein sequences obtained were retrieved for further analysis and the rest having significant similarities with the host were excluded.

### Essential Protein Screening

Accordingly, the essential proteins among the non-homologous proteins were identified utilizing a BLASTp search (with a threshold e value of 10^-100^) against the Database of Essential Genes and protein sequences demonstrating notable similarity with the DEG database were categorized as being vital for the survival of the pathogen and were selected for subsequent analysis [44–47].

### Metabolic Pathway Exploration

The association of the chosen essential proteins in diverse metabolic pathways of the host and the pathogen was achieved by KEGG automated annotation server (KAAS)[48]. The three letter organism codes ‘sag’, ‘san’, and ‘sak’ for *S. agalactiae* and ‘hsa’ for *H. sapiens* were selected, and employing the BHH method in the KAAS server, the metabolic pathways were collected independently and compared.

### Subcellular Localization Prediction

Subcellular location of the chosen essential proteins was initially performed by the pSORTbv3.03 tool [49,50] and cross-verified by CELLOv2.5 [51,52]. Both tools provide accurate predictions for various subcellular locations, encompassing proteins found in the cytoplasm, cytoplasmic membrane, extracellular space, cell wall, and those with unknown origins [49–52].

### Virulent Protein Screening

Accordingly, virulent proteins from the shortlisted essential proteins were retrieved from the tool VirulentPred2.0 [53]. The tool utilizes up-to-date datasets and advanced machine learning techniques to forecast the virulence status of proteins based on their PSSM profile [53].

### Determination of Physiochemical Parameters

The ProtParam server of ExPaSy was used to determine the physicochemical properties of the selected proteins [54].

### Analysis of Protein Interaction Networks

The potential interactions of the target proteins, with other proteins in *S. agalactiae* were predicted by the STRING database [55]. To prevent false-positive results, protein-protein interactions (PPI) studies were only considered if they had a high confidence score of 70% (0.700), choosing proteins with close interactions with at least three other proteins for further investigation.

### Prediction and Assessment of Three-Dimensional Structures

The three-dimensional structures of the target proteins were initially generated by the Swiss-Model online tool [56]. The tool uses homology-based approaches to construct 3D structures of proteins from amino acid sequences [56]. The 3D models of the selected proteins were constructed from template models that exhibited the most Global Mean Quality Estimate (GMQE) score and identity percentage [57–59] and was further validated by Swiss-Model structure validation [60,61].

## Results

Our study aimed to identify potential drug targets to combat the *S. agalactiae* serotype V strain, meeting the efficacy criteria of drug targets. The efficacy criteria include the targets must be non-homologous to humans, essential for the organism, and integral to the microbe’s main metabolic processes. The subtractive genomic analysis process is briefly presented in **Table 1**.

**Table 1.** Brief presentation of the subtractive genomic analysis of *S. agalactiae* serotype V.

| SI No. | Subtractive Approaches | Bioinformatics Tools and Servers Utilized | Number of Proteins |
| --- | --- | --- | --- |
| 1 | The whole proteome of *S. agalactiae* serotype V<br>(Strain ATCC BAA-611 / 2603 V/R) | UniProt | 2105 |
| 2 | Peer-Reviewed Proteins of *S. agalactiae* serotype V<br>(Strain ATCC BAA-611 / 2603 V/R) | UniProt | 390 |
| 3 | Proteins nonhomologous to<br>*H. sapiens* | BLASTp (E-value $10^{-4}$) | 200 |
| 4 | Essential proteins | DEG server (E-value $\leq 10^{-100}$) | 68 |
| 5 | Essential Proteins involved<br>only in unique metabolic<br>pathways | KAAS at KEGG | 51 |
| 6 | Proteins assigned KO (KEGG<br>Orthology) but not in any<br>pathway | KEGG Orthology | 15 |
| 7 | Essential membrane proteins | PSORTb | 12 |
| 8 | Essential membrane proteins | CELLO | 8 |
| 9 | Essential virulent proteins | VirulentPred2.0 | 6 |
| 10 | Essential membrane and virulent proteins | PSORTb,<br>CELLO,<br>VirulentPred2.0 | 2 |

### Protein Sequence Retrieval

The total proteins available in the reference proteome of this strain in UniProt was 2105, among which 390 protein sequences were found to be Swiss-Prot (Peer-reviewed) proteins. For further analysis, all peer-reviewed proteins were collected in FASTA format, while excluding the non-reviewed proteins.

### Finding Non-homologous Proteins

To identify non-homologous proteins, a BLASTp search of the NCBI database (with e-value threshold of 10^−4^) was conducted against *Homo sapiens* on 390 peer-reviewed proteins. Out of the 390 sequences, 200 were identified as non-homologous proteins, while the rest had resemblance to humans. While homologous protein sequences were excluded, these 200 non-homologous proteins were chosen for further analysis.

### Finding Essential Proteins

To identify essential proteins of the pathogen, a BLASTp search against the database of DEG (with an e-value cut off at 10^-100^) was performed. A total of 68 proteins (**S1 Table**) out of 200 were identified as crucial to the bacterium’s survival and were chosen for subsequent steps.

### Metabolic Pathway Exploration

The KAAS server revealed that 51 out of 68 essential proteins are associated with 41 distinct metabolic pathways, with no host pathways involved. Moreover, no common pathway between the host and the pathogen was observed, leading to the conclusion that the selected non-homologous proteins showed no involvement in the host pathways. Moreover, all 68 essential proteins had KEGG Orthology (KO) identifiers (**S2 Table**) by the KAAS server at KEGG, except for two proteins (namely, Bis(5’-nucleosyl)-tetraphosphatase (symmetrical) and galactosyl transferase CpsE). The unique metabolic pathways are listed in **Table 2**.

**Table 2.** Metabolic pathways unique to *S. agalactiae* serotype V.

| Sl. No | Unique Pathways | Total Proteins | Pathway ID | Proteins Involved |
| --- | --- | --- | --- | --- |
| 1 | Pentose phosphate pathway | 1 | 00030 | DEOB_STRA5 |
| 2 | Amino sugar and nucleotide sugar metabolism | 2 | 00520 | MURA_STRA5, MURB_STRA5 |
| 3 | Pyruvate metabolism | 2 | 00620 | ACKA_STRA5, CAPP_STRA5 |
| 4 | Propanoate metabolism | 1 | 00640 | ACKA_STRA5 |
| 5 | Oxidative phosphorylation | 2 | 00190 | ATPD_STRA5, PPAC_STRA5 |
| 6 | Photosynthesis | 1 | 00195 | ATPD_STRA5 |
| 7 | Carbon fixation by Calvin cycle | 1 | 00710 | CAPP_STRA5 |
| 8 | Other carbon fixation pathways | 2 | 00720 | CAPP_STRA5, ACKA_STRA5 |
| 9 | Methane metabolism | 2 | 00680 | CAPP_STRA5, ACKA_STRA5 |
| 10 | Fatty acid biosynthesis | 2 | 00061 | FABH_STRA5, FABZ_STRA5 |
| 11 | Glycerolipid metabolism | 2 | 00561 | PLSX_STRA5, PLSY_STRA5 |
| 12 | Glycerophospholipid metabolism | 1 | 00564 | PLSY_STRA5 |
| 13 | Purine metabolism | 1 | 00230 | DEOB_STRA5 |
| 14 | Pyrimidine metabolism | 3 | 00240 | PYRH_STRA5, KCY_STRA5, KTHY_STRA5 |
| 15 | Glycine, serine and threonine metabolism | 1 | 00260 | KHSE_STRA5 |
| 16 | Cysteine and methionine metabolism | 1 | 00270 | KHSE_STRA5 |
| 17 | Phenylalanine, tyrosine and tryptophan biosynthesis | 4 | 00400 | AROB_STRA5, AROD_STRA5, AROA_STRA5, AROC_STRA5 |
| 18 | Taurine and hypotaurine metabolism | 1 | 00430 | ACKA_STRA5 |
| 19 | D-Amino acid metabolism | 4 | 00470 | ALR_STRA5, DDL_STRA5,<br>MURI_STRA5, MURD_STRA5 |
| 20 | Peptidoglycan biosynthesis | 7 | 00550 | MURA_STRA5, MURB_STRA5,<br>MURC_STRA5, MURD_STRA5,<br>DDL_STRA5, UPPP_STRA5,<br>MURG_STRA5 |
| 21 | Teichoic acid biosynthesis | 1 | 00552 | UPPP_STRA5 |
| 22 | Thiamine metabolism | 1 | 00730 | RSGA_STRA5 |
| 23 | Nicotinate and nicotinamide<br>metabolism | 1 | 00760 | NADE_STRA5 |
| 24 | Pantothenate and CoA biosynthesis | 1 | 00770 | COAD_STRA5 |
| 25 | Biotin metabolism | 1 | 00780 | FABZ_STRA5 |
| 26 | RNA polymerase | 1 | 03020 | RPOA_STRA5 |
| 27 | Ribosome | 3 | 03010 | RS3_STRA5, RS4_STRA5,<br>RL10_STRA5 |
| 28 | Aminoacyl-tRNA biosynthesis | 2 | 00970 | SYGA_STRA5, SYGB_STRA5 |
| 29 | Protein export | 1 | 03060 | SECA_STRA5 |
| 30 | RNA degradation | 1 | 03018 | RNY_STRA5 |
| 31 | DNA replication | 1 | 03030 | RNH3_STRA5 |
| 32 | Nucleotide excision repair | 1 | 03420 | UVRC_STRA5 |
| 33 | Mismatch repair | 1 | 03430 | EX7L_STRA5 |
| 34 | Homologous recombination | 5 | 03440 | RECF_STRA5, RECO_STRA5,<br>RECR_STRA5, RECA_STRA5,<br>RUVA_STRA5 |
| 35 | ABC transporters | 1 | 02010 | PSTS1_STRA5 |
| 36 | Bacterial secretion system | 1 | 03070 | SECA_STRA5 |
| 37 | Two-component system | 3 | 02020 | PSTS1_STRA5, DNAA_STRA5,<br>LYTS_STRA5 |
| 38 | Cell cycle - Caulobacter | 2 | 04112 | DNAA_STRA5, MURG_STRA5 |
| 39 | Quorum sensing | 1 | 02024 | SECA_STRA5 |
| 40 | Tuberculosis | 1 | 05152 | PSTS1_STRA5 |
| 41 | Vancomycin resistance | 3 | 01502 | DDL_STRA5, ALR_STRA5,<br>MURG_STRA5 |

### Subcellular Localization Prediction

The pSORTbv3.0.3 tool predicted eight cytoplasmic membrane proteins, 51 cytoplasmic proteins, and nine proteins with uncertain or unknown subcellular localization from the shortlisted essential proteins. Subsequently, CELLOv2.5 predicted 12 membrane proteins, 54 proteins as cytoplasmic, and two extracellular proteins. However, both CELLOv2.5 as well as pSORTbv3.03 did not exhibit any proteins belonging in the cell wall region (**S3 Table**). The predicted results of the subcellular localization of essential proteins of *S. agalactiae* are presented in **Fig 3**. Out of the 8 and 12 cytoplasmic membrane proteins predicted by pSORTbv3.03 and CELLOv2.5, respectively, five proteins were predicted by both tools (**Table 3**).

**Fig 2.**
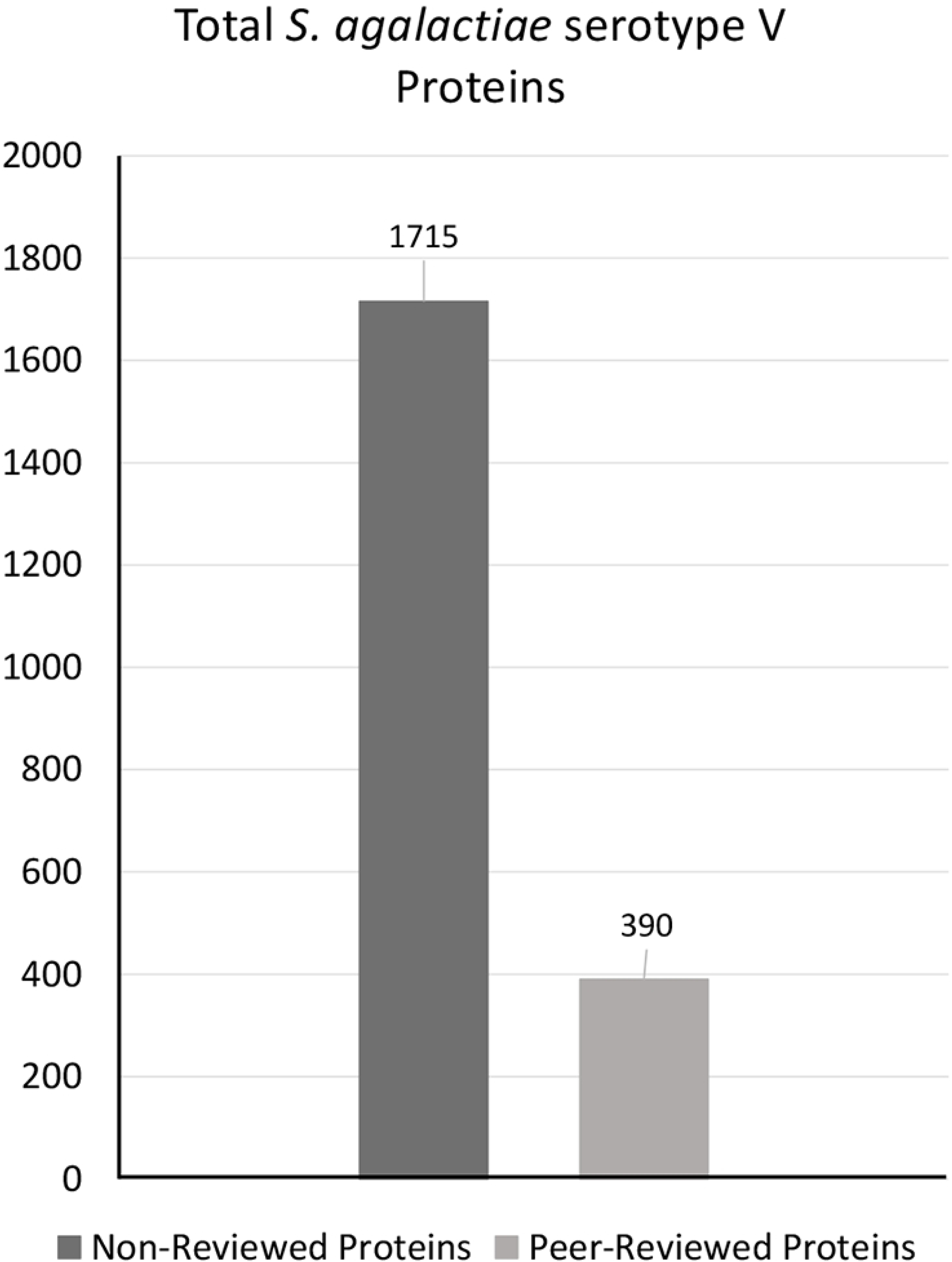
Total proteins of *S. agalactiae* serotype V. This shows the protein counts of non-reviewed and selected peer-reviewed sequences.

**Fig 3.**
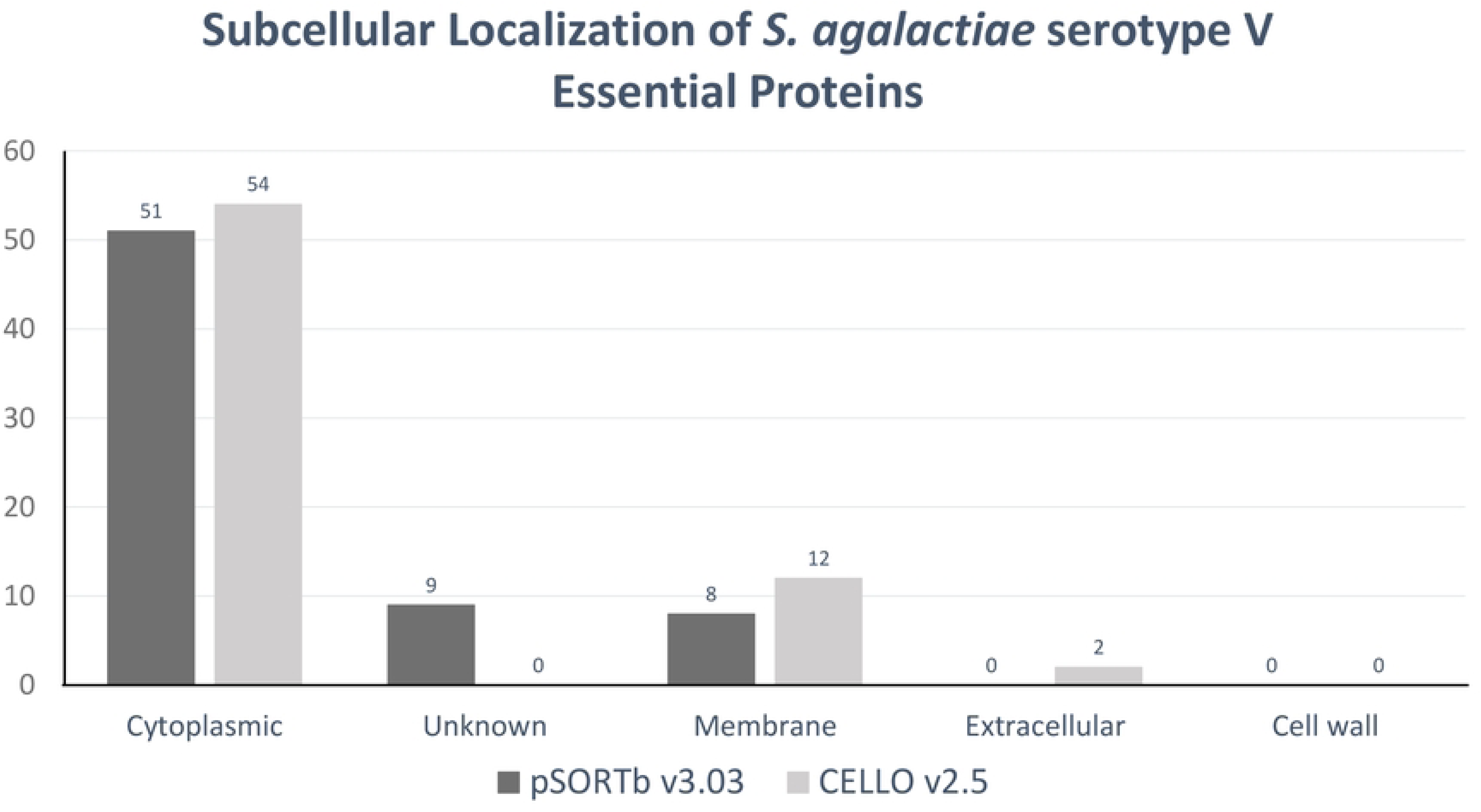
Essential, non-homologous proteins and their subcellular localization by pSORTbv3.03 and CELLOv2.5. This shows the protein counts of sequences predicted by subcellular localization prediction.

**Table 3.**
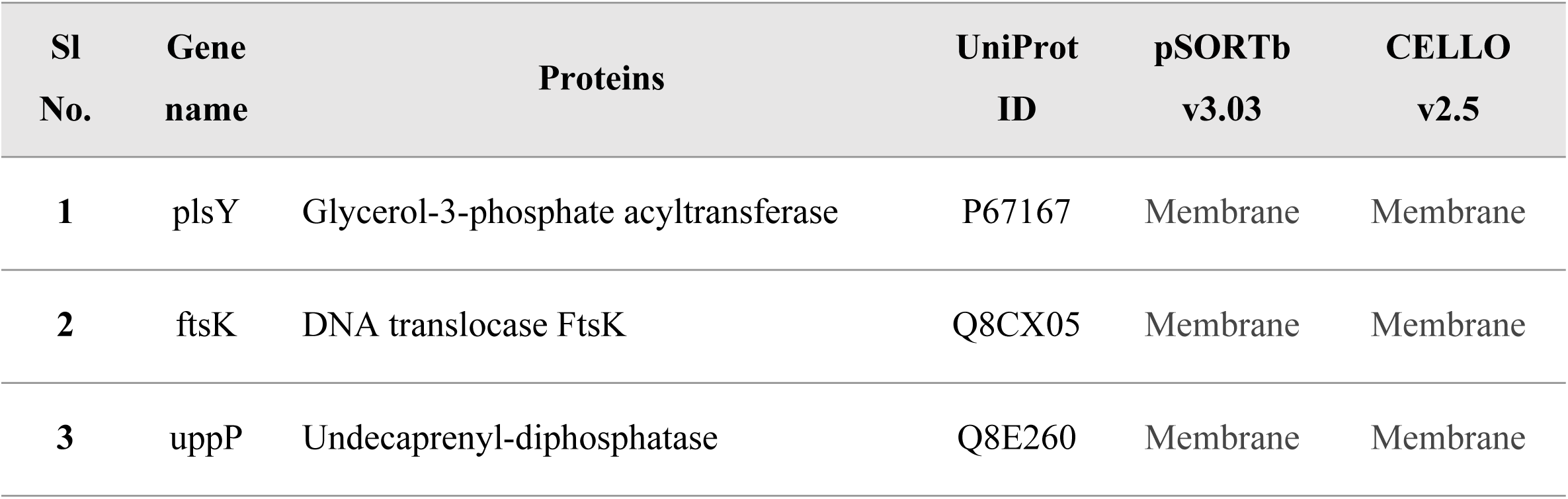
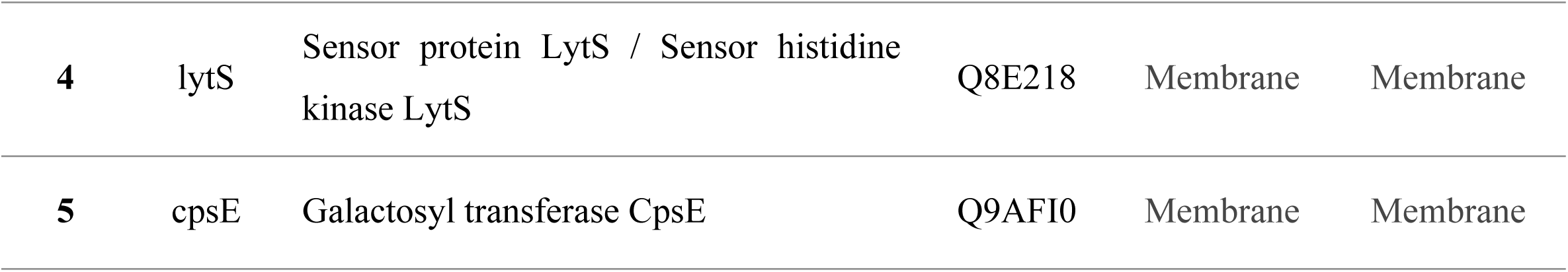
List of cytoplasmic membrane proteins identified by both pSORTbv3.03 and CELLOv2.5.

### Prediction of Virulent Proteins

The identification of virulent proteins, among 68 essential non homologous proteins, using VirulentPred2.0 revealed six proteins that are listed in **Table 4**. Inhibition of these proteins could significantly impact the pathogen’s functionality within the host organism. Among the six predicted proteins, LytS and CpsE were found to be both virulent and present in cytoplasmic membrane which implies that these two proteins are potential drug targets **Table 4** (Bold and Italic).

**Table 4.** Essential, non-homologous proteins predicted to be virulent by VirulentPred2.0.

| Sl No. | Gene name | Proteins | UniProt ID | KO ID | VirulentPred2.0 |
| --- | --- | --- | --- | --- | --- |
| 1 | hslO | 33 kDa chaperonin | P64402 | K04083 | Virulent |
| 2 | ruvA | Holliday junction branch migration complex subunit RuvA | Q8DWW6 | K03550 | Virulent |
| 3 | pstS | Phosphate-binding protein PstS | Q8DZV4 | K02040 | Virulent |
| 4 | *lytS* | *Sensor protein LytS* | Q8E218 | K07704 | Virulent |
| 5 | recO | DNA repair protein RecO | Q8E2G8 | K03584 | Virulent |
| 6 | *cpsE* | *Galactosyl transferase CpsE* | Q9AFI0 | - | Virulent |

### Physiochemical Property Analysis

The physiochemical properties of the selected two proteins, namely Sensor protein LytS and Galactosyl transferase CpsE, were analyzed by ProtParam tool. The analysis showed that both proteins are slightly hydrophobic, basic, stable, and highly thermostable in nature. It is important to note that Galactosyl transferase CpsE is more basic in nature than Sensor Protein LytS. The detailed results of physiochemical property analysis of two proteins are presented in **Table 5**.

**Table 5.** ProtParam analysis of Sensor protein LytS and Galactosyl transferase CpsE.

| Physicochemical Properties | Sensor protein<br>LytS | Galactosyl transferase<br>CpsE |
| --- | --- | --- |
| Amino acids | 581 | 449 |
| Molecular weight | 64548.62 | 52364.13 |
| Theoretical pI | 7.71 | 9.34 |
| Instability Index | 35.59 | 33.46 |
| Aliphatic Index | 107.01 | 103.94 |
| Grand average of hydropathicity (GRAVY) | 0.075 | 0.059 |
| Estimated half-life in Mammalian reticulocytes, in-vitro | 30 hours | 30 hours |
| Estimated half-life in Yeast in-vivo | >20 hours | >20 hours |
| Estimated half-life in *Escherichia coli*, in-vivo | >10 hours | >10 hours |

### Analysis of Protein Interaction Networks

The significant association of these two chosen proteins with other proteins in the pathogen was identified using the STRING database, and the amino acid sequences of Sensor protein LytS and Galactosyl transferase CpsE was uploaded to the server. The Sensor protein LytS developed 4 PPI networks (**Fig 4A**) depicted as lytS in red node. LytS had 4 edges, 3 anticipated edges, 4 nodes, with a 2.0 average node degree. Protein-Protein Interaction enrichment p-value was 0.331, with 0.833 average local clustering coefficient. Moreover, it interacted with two neighbouring two-component system response regulator proteins (lytR and SAG1016) and one neighbouring conserved hypothetical protein (SAG0184). On the other hand, Galactosyl transferase CpsE had 20 PPI networks (**Fig 4B**) represented as cpsE in red node. There were around 20 nodes, 125 edges, with an average node degree of 12.5, a clustering coefficient of 0.888, 21 predicted edges, and a PPI enrichment p-value of more than 1.0e-16. Furthermore, it interacted with numerous neighbouring proteins mostly involved in the assembly, regulation and biosynthesis of the GBS capsular polysaccharide (cpsD, cpsC, cpsB, cpsL, cpsA, cpsJ, cpsG, cpsH, cpsM, cpsO, cpsN, cpsK, cpsF) along with different other neighbouring important enzymes (neuC, neuD, neuA, neuB, etc).

**Fig 4.**
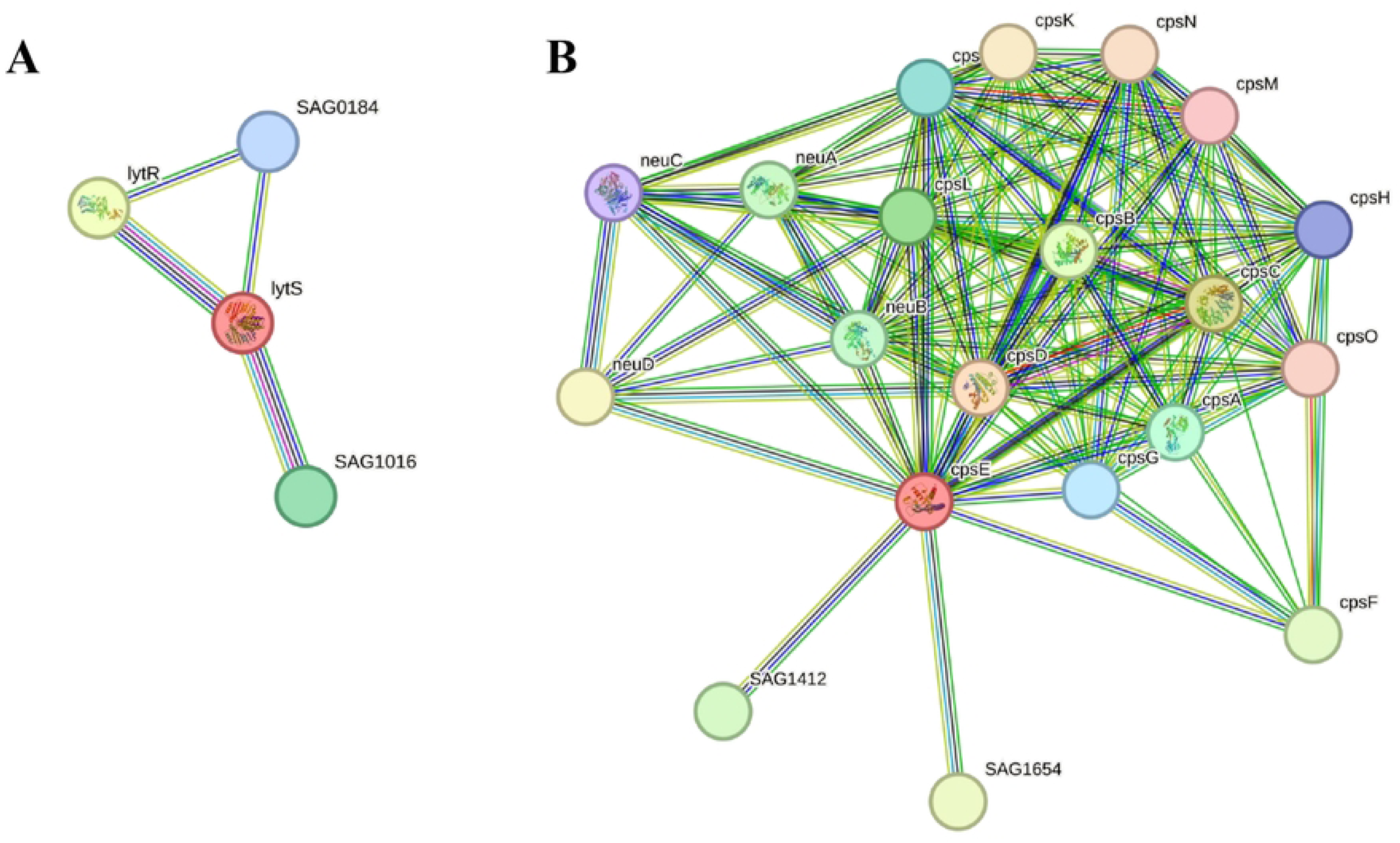
Prediction of protein-protein interactions of the two proteins. The chosen proteins are represented in red nodes. Each node represents all the proteins produced by a single, protein-coding gene locus. Empty nodes represent proteins of unknown 3D structure and filled nodes indicates that a 3D structure is known or predicted. Edges represent protein-protein associations. (A) Sensor protein LytS protein. (B) Galactosyl transferase CpsE protein.

### Prediction and Assessment of Three-dimensional Structures

The online server Swiss-Model generated fifty initial templates for each protein, and the templates with the most GMQE score and identity percentage were chosen to construct the 3D models. Moreover, the predicted structures of Sensor protein LytS and Galactosyl transferase CpsE had 0.80 and 1.06 MolProbity scores with 97.58% and 95.97% of residues in the favored regions of the Ramachandran plot respectively, according to the Swiss-Model structure validation (**Fig 5)**.

**Fig 5.**
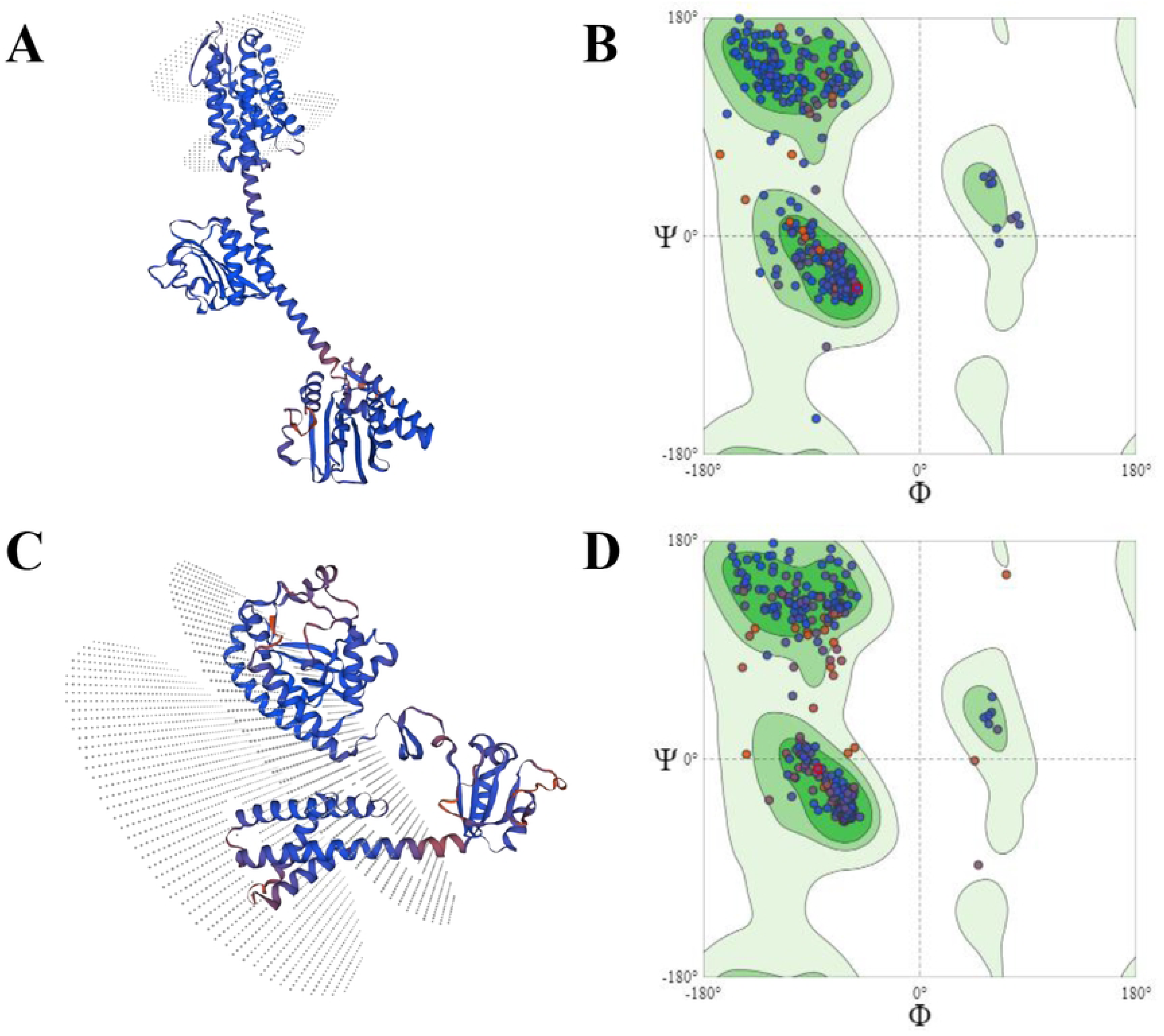
Determination of three-dimensional structure. (A) Three-dimensional structure of Sensor protein LytS protein. (C) Three-dimensional structure of Galactosyl transferase CpsE protein. (B) The Ramachandran Plot of Sensor protein LytS protein. (D) The Ramachandran Plot of Galactosyl transferase CpsE protein.

## Discussion

Developing new therapeutic drugs and vaccines are challenging. Advancements in computational research, sequence-based technologies, and the availability of diverse pathogens’ genomes and proteomics data have made it easier to perform the task. *In silico* subtractive genome techniques are promising in identifying specific genes and proteins in various organisms including *Clostridium botulinum* [62]*, Mycoplasma pneumoniae* [31]*, Streptococcus pneumonia* [63], *Campylobacter jejuni* [64]*, Salmonella typhi* [26], *Legionella pneumophila* [39], *Arabidopsis thaliana* [65], *Meningococcus B* [66]*, Eubacterium notadum* [67], *Fusobacterium nucleatum* [68], *Salmonella enterica subsp. Poona* [69], *Treponema pallidum* [70], *Staphylococcus aureus N315* [71], *Acinetobacter baumanii* [72], *Bartonella bacilliformis* [73], *Bordetella pertussis* [74], *Serratia marcescens* [75], and *Staphylococcus aureus* [76].

In this study, we employed a subtractive genomics-based computational approach to screen the entire proteome of *S. agalactiae* serotype V (ATCC BAA-611 / 2603 V/R) for the identification of potential drug targets. We obtained the entire proteome from UniProt database and selected only 390 peer-reviewed sequences for subsequent investigation. The study chose exclusively peer-reviewed sequences to avoid overrepresentation of specific proteins and computational difficulties, ensuring the accuracy and representativeness of the resulting data, thereby reducing redundancy within the proteome. The database of UniProtKB and its proteome portal houses over 227 million sequence information and 451,000 collections of proteomes respectively, derived from fully sequenced archaeal, bacterial, eukaryotic and viral genomes [40–42].

Afterwards, the BLASTp analysis against the host *H. sapiens* revealed 200 non-homologous proteins to the host. Proteins that participate in various fundamental cellular systems have been identified as homologous, exhibiting analogous functions in both humans and bacteria throughout the course of evolution [77,78]. We excluded the remaining proteins because cytotoxic reactions and adverse effects might arise from drug targets that are similar to the host genome and the aim of targeting such homologous proteins might result in detrimental consequences [79,80]. In some studies, human homologous proteins (up to 40% similarity) were selected to identify potential drug targets [81], based on the idea that low similarity in sequence would produce a slightly different protein structure from human host and it would be safely considered as a drug target [81,82]. This could be the case, when no potential drug targets are not found in non-homologous sequence pool.

Later on, we utilized the DEG database, that successfully lead to the selection of 68 essential non-similar proteins out of 200 non-homologous sequences (**S1 Table**). The protein sequences that exhibited significant homogeneity with the DEG database were classified as essential for the survival of the bacterium. The promising candidates for organism-specific drug targets are proteins that are vital for the pathogen’s survival and do not share homology with the host genome [47,72,73,80,83].

Subsequently, KEGG KAAS analysis revealed 51 essential non-homologous proteins involved in 41 unique pathways (**Table 2**). These unique pathways including two-component system, quorum sensing, peptidoglycan biosynthesis, and various amino acid metabolism systems such as alanine, aspartate, glutamate, cysteine, methionine, tyrosine, histidine metabolism, etc., are essential for the survival of *S. agalactiae* and when interrupted would cause the bacterium to not function properly [30,64]. Environmental stress can increase gene expression and metabolic processes in bacteria, potentially leading to the development of resistant pathogens with distinct metabolic processes [38,76]. Thus, these essential proteins involved in such unique pathways alone could be excellent targets for drugs and vaccines [38,76].

In addition, protein localization is crucial in drug development, as it governs the formulation and design of novel drugs and vaccines, as demonstrated in studies including *Acinetobacter baumanii* [22,72], *Bartonella bacilliformis* [73], *Bordetella pertussis* [74], *Streptococcus pneumoniae* [33,84], *Salmonella typhi* [24], *Mycoplasma genitalium* [85], *Neisseria meningitides* [36], *Staphylococcus aureus* [38,76], and *Treponema pallidum* [70]. In several studies, cytoplasmic proteins were selected as drug targets for their availability and involvement in metabolic pathways crucial to the survival of those pathogens [20,34,85,86]. However, in our study we emphasized on the cytoplasmic membrane proteins in view of the fact that they can further be utilized as a target for vaccine development, as membrane proteins are considered to be the best criteria for vaccines [63,87,88]. Moreover, we observed that the selected cytoplasmic membrane proteins were involved in essential two component system pathway and the GBS polysaccharide formation pathway of *S. agalactiae* and had virulent characteristics. Virulent proteins in pathogens play a crucial role in bacterial pathogenesis along with the degradation and control of the host immune system mechanisms [89,90]. Moreover, it was seen that membrane proteins are essential for integral cellular signal detection, transduction, and various other biological processes [91]. They act as diffusion barriers for ions, water, transport systems, and nutrients and when disrupted or degraded, can lead the bacteria to cease functioning and can serve as excellent druggable targets [91]. It is important to note that a significant number of druggable targets were derived from membrane proteins [21,87,88,91–96].

The pSORTb v3.03 tool is widely regarded as the most effective approach for predicting subcellular localization, with a classification precision measured around 96%[49,50] and therefore it was initially utilized to predict the subcellular localization of the chosen essential proteins. Subsequently, CELLOv2.5 tool was employed for the cross-validation of the chosen proteins **(S3 Table).** As a result, 8 proteins from pSORTbv3.03, and 12 proteins from CELLOv2.5 were predicted to be cytoplasmic membrane proteins. Moreover, two proteins from pSORTbv3.03, six proteins from CELLOv2.5 were revealed to be cytoplasmic membrane proteins and among them five were found to be common (**Table 3**) which were used in the subsequent analysis.

Furthermore, virulent factors in pathogens facilitate bacterial adhesion, colonization, invasion, and disease pathogenesis [89,90]. The identification of unique virulent factors of *S. agalactiae* would represent a substantial contribution, given that these factors are crucial in the control or degradation of the host immune system [97]. Hence, from the set of essential proteins, six virulent proteins (**Table 4**) were predicted by the tool VirulentPred2.0, implying that inhibition of these proteins could render the pathogen non-virulent and significantly impact the pathogen’s functionality within the host organism [23,25,26,35,62,68]. The two proteins, namely sensor protein LytS and galactosyl transferase CpsE, had the potential to be utilized as promising drug candidates because of their virulent nature as well as cytoplasmic membrane characteristics and were selected for further analysis.

The STRING server afterwards revealed that the two selected proteins could function as core proteins associating with three or more neighboring proteins of *S. agalactiae*. As a result, repressing these proteins can inhibit the proper functioning of other related proteins [32,34,39,75]. It was found that Sensor protein LytS, also known as sensor histidine kinase LytS, encoded by the gene lytS, is a part of the Two-component Regulatory System LytR/LytS, and played a pivotal role in the regulation of cell wall metabolism. The system which enables bacteria to perceive, respond, and cope with stressful conditions by gene expression regulation is the two-component signal transduction system [32]. The sensor histidine kinases are capable of detecting alterations in the pathogen’s external environment and transmitting signals that induce modifications in the internal machinery of bacterial cells, enabling them to adapt and exploit these changes [34]. ProtParam analysis predicted that it is hydrophobic, basic, stable, and thermostable in nature with 581 amino acids in length (**Table 5**). Moreover, by observing its neighboring proteins (**Fig 4A**), it was discovered that the sensor protein LytS interacted with two neighboring two-component system proteins which were response regulator proteins and one conserved hypothetical protein. Conversely, Galactosyl transferase CpsE, encoded by the gene cpsE, was observed in the biosynthesis of capsular polysaccharide and played a pivotal role in the assemblage of capsular polysaccharide in group B streptococci. The polysaccharide capsule represents a significant virulence factor in pathogens belonging to the *Streptococcus* genus and played a significant role in enabling these bacteria to evade the innate immune response by providing protection against phagocytosis, opsonization, along with the complement system [98]. ProtParam analysis predicted that it was hydrophobic, basic, stable and thermostable in nature with 449 amino acids in length (**Table 5**). Moreover, by observing its neighboring proteins (**Fig 4B**), it was discovered that the galactosyl transferase CpsE interacted with numerous proteins mostly involved in the assembly, regulation and biosynthesis of the GBS capsular polysaccharide along with different other important enzymes.

In addition, SWISS-MODEL server effectively predicted, analyzed, and evaluated the 3D structures of the two chosen proteins (**Fig 5**). Predicting protein three-dimensional structures significantly aids in understanding protein dynamics, functions, ligand interactions, and other protein components [99]. It was seen that the residues in the Ramachandran favoured regions of both proteins were more than 85% and the Molprobity score was between the expected range of −4 to 2 [100], thereby confirming that the above values and calculations satisfied the structural validation criteria and showed that the predicted structures were of high quality as reported in various other studies [20,25,26,32,34,35,37,39,65,67,69,70,75,76,101,102].

As a result, we successfully identified and analyzed new proteins that show great promise as potential therapeutic targets. The potential of these proteins was prudently determined pertaining to their fundamental contribution to play an essential part in the survival of the pathogen and their feasibility in combating *S. agalactiae* serotype V (Strain ATCC BAA- 611/2603V/R). As far as we are aware, this serotype has never been the subject of a subtractive genomics study before, and we believe it will provide a promising new strategy for preventing the spread of the antibiotic-resistant bacteria. Furthermore, we have investigated the druggability of these two proteins on other pathogenic bacteria including *S. agalactiae* serotype III, *Shigella flexneri, Streptococcus pneumoniae, Klebsiella pneumoniae, Acinotobacter baumanii, Enterococcus faecium, Pseudomonas aeruginosa, Shigella sonnei, Shigella dysenteriae, Bacillus cereus, Bacillus anthracis, Staphylococcus aureus, Clostridium perfringens, Mycobacterium tuberculosis, Salmonella Typhi, Yersinia enterocolitica, Campylobacter jejuni, Listeria monocytogenes, Vibrio cholerae,* and *Clostridium botulinum,* and found these two proteins have almost 70-100% similarity with different sensor histidine kinases/sensor proteins and sugar transferases of other pathogenic bacteria and are involved in various essential cellular processes (Data not shown). However, it requires further investigations which is not in the scope of the current article.

## Conclusion

The development of new drugs has been significantly expedited by the utilization of bioinformatics tools to extract and analyze genome and proteome sequences from diverse pathogens across multiple databases. The significance of utilizing *in silico* methods for discovering novel therapeutic targets having least homology to the host proteome is monumental. The subtractive genomics methodology is impressively capable of effectively resolving challenges which are prevalent in traditional drug discovery methodologies. Using subtractive genomics approach, we have successfully identified two proteins, namely LytS and CpsE, that have the potential to be used as drug targets. It is reasonable to assume that the identification of these two druggable proteins in *S. agalactiae* serotype V (Strain ATCC BAA-611/2603V/R) will lay the foundations for developing novel therapeutic drugs for associated infections.

## Supporting Information

**S1 Table**

**S2 Table**

**S3 Table**

